# The CIP2A-TOPBP1 complex safeguards chromosomal stability during mitosis

**DOI:** 10.1101/2021.02.08.430274

**Authors:** Mara De Marco Zompit, Clémence Mooser, Salomé Adam, Silvia Emma Rossi, Alain Jeanrenaud, Pia-Amata Leimbacher, Daniel Fink, Daniel Durocher, Manuel Stucki

## Abstract

The accurate repair of DNA double-strand breaks (DSBs), highly toxic DNA lesions, is crucial for genome integrity and is tightly regulated during the cell cycle. In mitosis, cells inactivate DSB repair in favor of a tethering mechanism that stabilizes broken chromosomes until they are repaired in the subsequent cell cycle phases. How this is achieved mechanistically is not yet understood, but the adaptor protein TOPBP1 is critically implicated in this process. Here, we identify CIP2A as a TOPBP1-interacting protein that regulates TOPBP1 localization specifically in mitosis. Cells lacking CIP2A display increased micronuclei formation, DSB repair defects and chromosomal instability. CIP2A is actively exported from the cell nucleus in interphase but, upon nuclear envelope breakdown at the onset of mitosis, gains access to chromatin where it forms a complex with MDC1 and TOPBP1 to promote TOPBP1 recruitment to sites of mitotic DSBs. Collectively, our data uncover CIP2A-TOPBP1 as a mitosis-specific genome maintenance complex.

---

Failure to accurately repair DSBs leads to genome instability, cell death or cancer^1^. Cells repair DSBs by two distinct families of pathways: non-homologous end joining (NHEJ) pathways that are the only DSB repair pathways active in G1, and homologous recombination (HR) pathways that require a sister chromatid as template and thus only occur in S and G2 phases^2^. By activating the G2/M checkpoint, cells buy time to repair the majority of DNA breaks before entering mitosis. However, DSBs can also occur during cell division, for example as a result of under-replicated DNA regions at common fragile sites, which impede chromosome segregation and may ultimately cause DNA breakage in mitosis^3–5^. Cells rewire their response to DSBs in mitosis, at least in part to prevent telomeric fusions^6^. There is evidence that, instead of repairing them, mitotic cells stabilize chromosomal breaks until they can be safely repaired in the subsequent cell cycle phases^6–9^. How this is achieved mechanistically is not yet understood, but what is clear is that the chromatin response to DSBs is truncated in mitosis^8^. Early events such as phosphorylation of H2AX and recruitment of MDC1 are intact, but further downstream responses that in interphase regulate DSB repair pathway choice are severed by the inability of mitotic cells to recruit RNF8/RNF168, 53BP1 and BRCA1^6,8^. Instead, MDC1 mediates the recruitment of TOPBP1 to mitotic DSB sites, a process that is critical for the maintenance of chromosomal stability^10^ TOPBP1 is a highly versatile adaptor protein implicated in several aspects of genome integrity maintenance^11^. It is composed of different domains and regions, including nine BRCT domains, and an ATR activation domain (AAD; see Fig. 1a). TOPBP1 undergoes prominent enrichment at DSB sites throughout the cell cycle by at least four distinct mechanisms that involve phosphorylation-dependent interactions of alternating combinations of its four phospho-binding BRCT domains (BRCT 1, 2, 5 and 7) with the adaptor proteins 53BP1, RAD9, Treacle and MDC1^12, 10,13,14^ TOPBP1 interacts with Casein kinase 2 (CK2) phosphorylated MDC1 via its N-terminal BRCT0-2 module. This interaction is not cell cycle regulated but paradoxically, MDC1 recruits TOPBP1 by direct interaction exclusively in mitosis^10^ Here we present a possible solution to this conundrum by identifying cancerous inhibitor of protein phosphatase 2A (CIP2A) as a TOPBP1-interacting protein that promotes TOPBP1 recruitment to sites of DSBs specifically in mitosis. We further demonstrate that loss of CIP2A leads to DSB repair defects, increased micronuclei formation and chromosomal instability. Finally, we propose that CIP2A-TOPBP1 complex formation is regulated during the cell cycle by nuclear export, which spatially sequesters CIP2A from TOPBP1 in interphase cells, thus allowing their efficient interaction exclusively in mitosis.

**Fig. 1.**
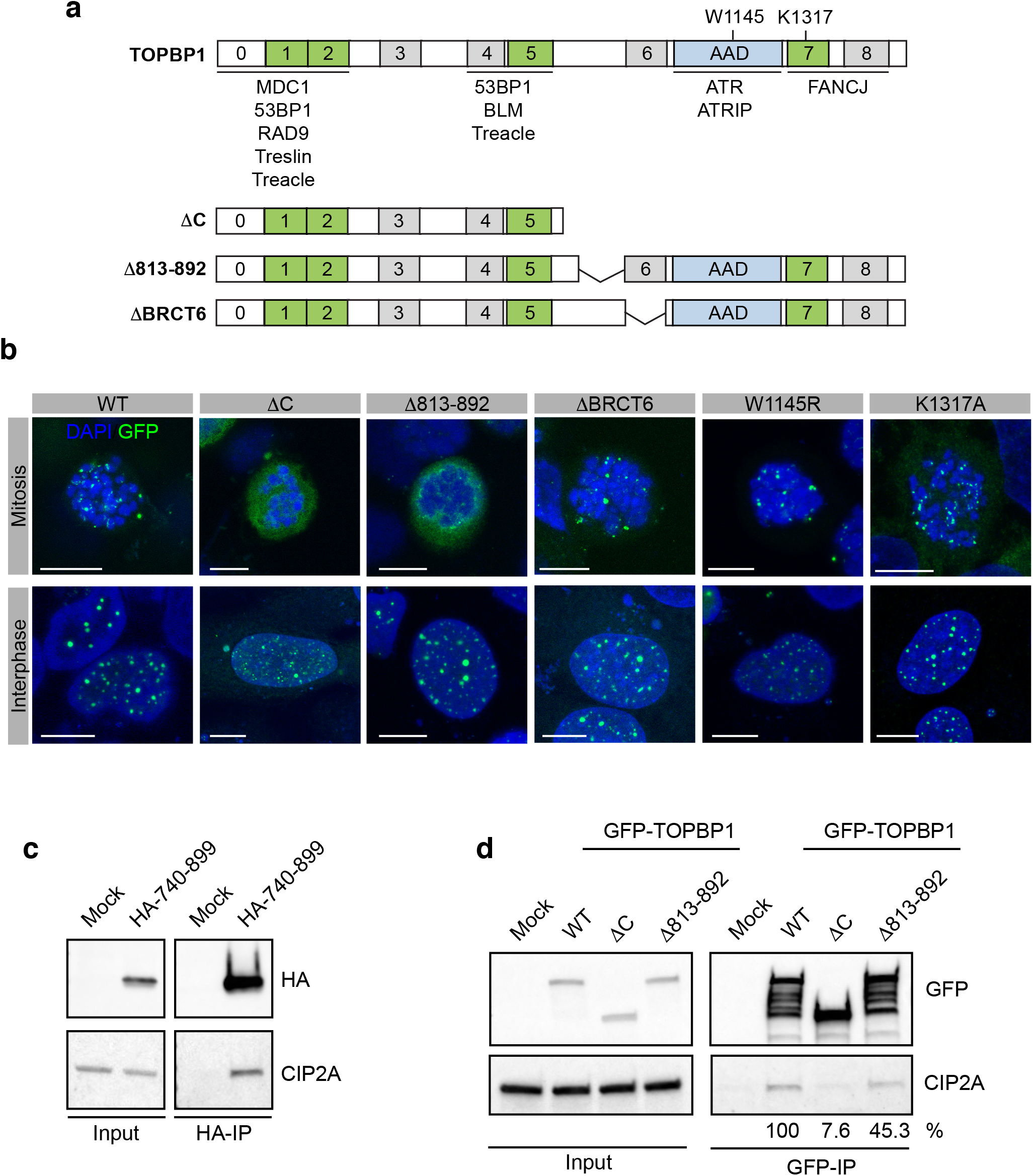
CIP2A is a TOPBP1 interaction partner. **a** Schematic showing the layout of conserved domains and regions in TOPBP1. Key amino acids in the AAD (W1145) and BRCT7 (K1317) are indicated. GFP-tagged full-length TOPBP deletion constructs of TOPBP1, lacking either the C-terminal portion of the protein (DC), a conserved region between BRCT5 and 6 (Δ813-892) and BRCT domain 6 (ΔBRCT6). **b** Localization of GFP-TOPBP1 wild type and mutants in mitotic cells after 1 Gy of IR and interphase cells after 3 Gy of IR. Displayed are maximum intensity projections of confocal z-stacks. All scalebars = 10 μm. **c** HA-immunoprecipitation from 293FT cells transfected with a HA-tagged TOPBP1 fragment spanning the entire region between BRCT5 and 6 (amino acids 740-899). **d** GFP-immunoprecipitation from 293FT cells transfected with GFP-tagged TOPBP1 wild type and indicated deletion mutants. Relative CIP2A band intensities are indicated.

## RESULTS

### CIP2A is a TOPBP1 interaction partner

We previously demonstrated that TOPBP1 is recruited to sites of DSBs in mitosis by direct phosphorylation-dependent interactions between its N-terminal BRCT domains with phosphorylated MDC1^10^. Interestingly, we observed that a C-terminal TOPBP1 deletion mutant containing BRCT domains 0-5 but lacking BRCT domains 6-8 and the AAD was recruited to sites of DSBs in G1, but not in mitosis (Fig. 1b). This suggested that domains/regions downstream of BRCT5 are important for TOPBP1 DSB recruitment in mitosis. Inactivating point mutations in BRCT7 and the AAD, and deletion of BRCT6 did not affect TOPBP1 recruitment, neither in interphase nor in mitosis (Fig. 1b). However, deletion of a conserved region between BRCT5 and 6 (amino acids 813-892) abrogated mitotic DSB recruitment, while recruitment in interphase was unaffected (Fig. 1a,b and Supplementary Fig. 1a). To test if this region constitutes a previously uncharacterized interaction surface for a protein that mediates TOPBP1 recruitment specifically in mitosis, we expressed a hemagglutinin (HA)-tagged protein fragment corresponding to the entire region between BRCT5 and 6 (human TOPBP1 residues 740-899) in 293-T cells, followed by immunoprecipitation with HA affinity beads. Bound proteins were identified by liquid chromatography-tandem mass spectrometry (LC-MS/MS). Among the 16 proteins present only in the HA-TOPBP1(740-899) pull-down but not in the control, CIP2A (also called KIAA1524) caught our attention (Supplementary Fig. 1b,c). We previously identified this protein in a proteomic screen for interaction partners of full-length TOPBP1^14^ Moreover, it was also identified in a recent proximity proteomic screen for BRCA1-interacting factors along with TOPBP1^15^. CIP2A was originally described as an endogenous inhibitory factor of protein phosphatase 2a (PP2A), and the protein is overexpressed in multiple cancers^16,17^. Interestingly, genotoxin sensitivity profiling by CRISPR-Cas9 drop-out screens revealed high sensitivity of CIP2A loss to ATR inhibitors and drugs that induce DSBs in proliferating cells such as Topoiomerase I and II inhibitors^18,19^. Inspection of one of these DNA damage chemogenetic datasets^19^ revealed a significant drug sensitivity correlation between MDC1 and CIP2A (Supplementary Fig. 1d). Combined, these were cues to further investigate a potential link between CIP2A and TOPBP1 in the mitotic DSB response. Western blotting confirmed that CIP2A was specifically pulled down by the conserved region between BRCT5 and 6 of TOPBP1 (Fig. 1c). In addition, co-immunoprecipitation with GFP-tagged full-length TOPBP1 and deletion mutants showed defective interaction with the C-terminal deletion mutant, and significantly reduced interaction upon deletion of amino acids 813-892 (Δ813-192; Fig. 1d). Together, these findings identify CIP2A as a TOPBP1 interacting factor that binds to a region in TOPBP1 that localizes between BRCT5 and 6.

### CIP2A interacts with TOPBP1 at sites of DSBs in mitosis

Consistent with a previous report^20^ we found that in undamaged cells, CIP2A is localizing to the cytoplasm in interphase and accumulates on centrosomes along with TOPBP1 in mitosis (Fig. 2a). Strikingly, upon irradiation, CIP2A was markedly enriched in foci on condensed mitotic chromosomes, where it co-localized with TOPBP1, while it was not detectable at sites of DSBs in interphase cells (Fig. 2b-d). By using high-resolution microscopy, we had previously observed that many IR-induced TOPBP1 structures in mitosis assumed the shape of filamentous assemblies^10^ Airyscan confocal microscopy showed that CIP2A was also present in such filaments on mitotic chromosomes and consistently co-localized with TOPBP1 (Fig. 2e). To test if CIP2A and TOPBP1 interact at sites of DSBs in mitosis we performed proximity ligation assays *(in situ* PLA). In unirradiated mitotic cells, the PLA signal was evenly distributed throughout the cell with occasional focal accumulation. Upon irradiation, the PLA signal was mostly concentrated in foci that colocalized with DAPI staining, indicating that CIP2A and TOPBP1 indeed interact at sites of DBSs on mitotic chromosomes (Fig. 2f). These data indicate that CIP2A is recruited to sites of DSBs exclusively in mitosis where it co-localizes with TOPBP1.

**Fig. 2.**
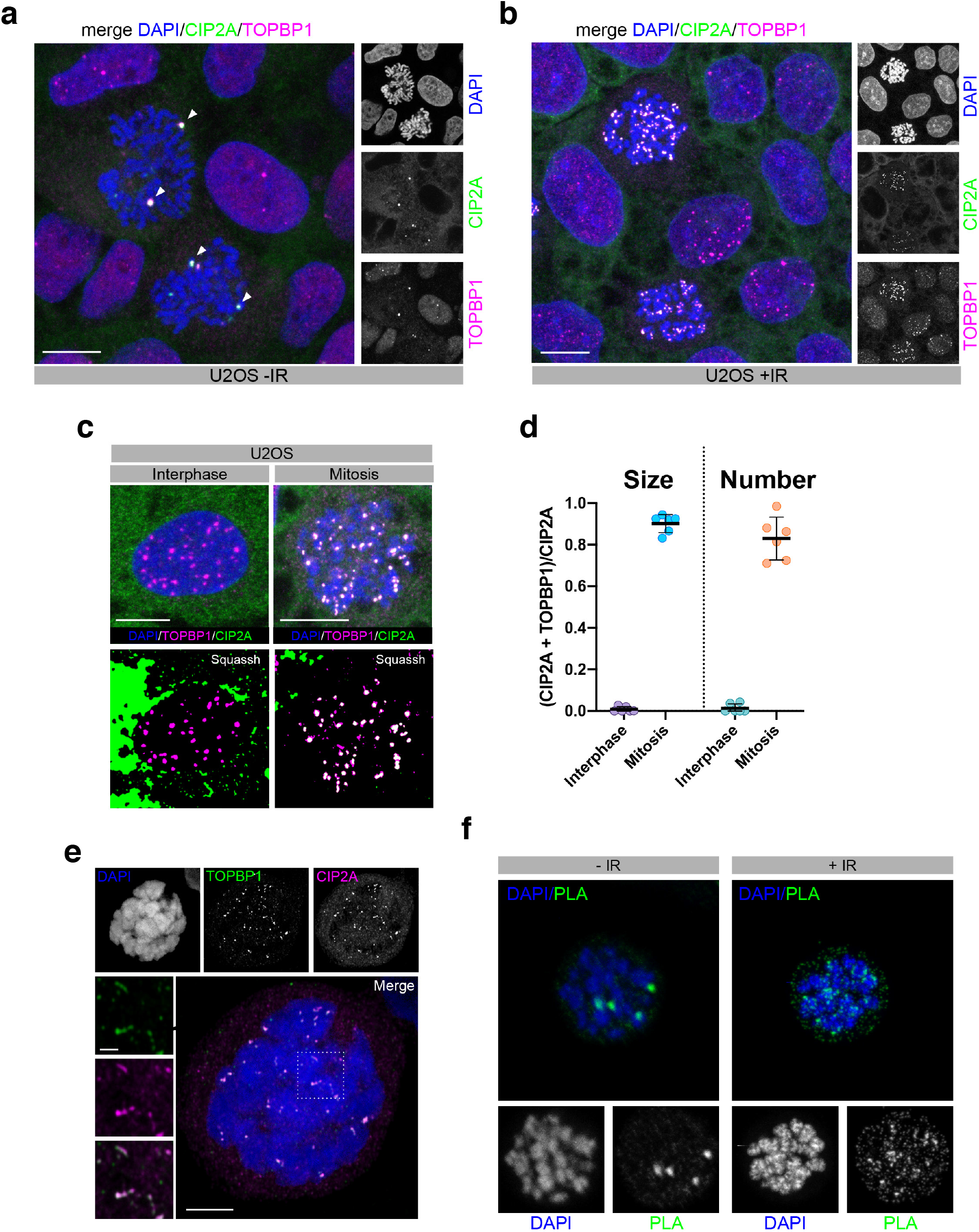
CIP2A interacts with TOPBP1 at sites of DSBs in mitosis. **a** Confocal micrograph (maximum intensity projection) of untreated U2OS cells, stained for TOPBP1 and CIP2A. Centrosomes are highlighted with white arrowheads. **b** Confocal micrograph (maximum intensity projection) of Nocodazole-arrested U2OS cells 1h after irradiation with 1 Gy, stained for TOPBP1 and CIP2A. **c** Upper panels: confocal micrographs of interphase U2OS cells treated with 3 Gy and U2OS cells arrested in mitosis by Nocodazole and treated with 1 Gy. Lower panels: micrographs deconvoluted and segmented by SQUASSH **d** Quantitative analysis of CIP2A and TOPBP1 colocalization by SQUASSH. Left: object size colocalization (area of object overlap divided by total object area). Right: object number colocalization (fraction of objects in each channel that overlap ≥ 50%). Each data point represents one cell (n=6). Bars and error bars represent mean and SD. **e** Airyscan high-resolution confocal image (maximum intensity projection) of CIP2A and TOPBP1 foci in mitosis 1 h after 1 Gy of IR. Scale bar in the merge panel: 5 μm; scale bar in the zoomed panels: 1 μm. **f** Detection of CIP2A-TOPBP1 co-localization by *in situ* PLA in U2OS cells arrested in mitosis by Nocodazole and mock treated or treated with 1 Gy of IR. All scale bars = 10 μm unless indicated otherwise.

### CIP2A mediates TOPBP1 DSB accumulation in mitosis

Next, we sought to determine if CIP2A is required for TOPBP1 recruitment to DSBs on mitotic chromosomes. To this end, we knocked out endogenous CIP2A in the hTERT immortalized non-transformed human retinal pigmented epithelial cell line RPE-1, using CRIPSR/Cas9 (henceforth termed ΔCIP2A; Fig. 3a and Supplementary Fig. 2a,b). TOPBP1 foci were completely absent in irradiated mitotic ΔCIP2A cells, but they were fully restored by stable transduction of ΔCIP2A cells with a lentiviral vector containing Flag-tagged wild type CIP2A cDNA (Fig. 3b,c), thus ruling out off-target effects of the guide RNA. TOPBP1 enrichment at mitotic DSBs was also completely defective in U2OS cells in which CIP2A was depleted by siRNA transfection, indicating that this effect is not cell type specific (Supplementary Fig. 2c). Depletion of TOPBP1 by siRNA also abrogated CIP2A foci in mitotic cells, suggesting that for efficient accumulation at sites of mitotic DSBs, TOPBP1 and CIP2A may be dependent upon each other (Fig. 3d). However, we also noted that depletion of TOPBP1 led to a marked reduction of CIP2A expression, but not vice versa, which could also have contributed to the apparent reduction of CIP2A foci in TOPBP1 depleted mitotic cells (Supplementary Fig. 2d). To test if TOPBP1 must exist in a complex with CIP2A in order to be recruited to sites of mitotic chromosome breaks, we knocked-down TOPBP1 expression in U2OS cells by 3’-UTR-targeting siRNA followed by re-expression of GFP-tagged wild type or TOPBP1 Δ813-192. CIP2A only accumulated in foci in the presence of wild type TOPBP1, but not in the presence of the Δ813-192 mutant, thus indicating that the recruitment of both of these proteins to sites of mitotic DNA breaks is dependent on their efficient interaction (Fig. 3e). We conclude from these results that CIP2A controls TOPBP1 recruitment to sites of mitotic DSBs and that CIP2A and TOPBP1 recruitment in mitosis are interdependent.

**Fig. 3.**
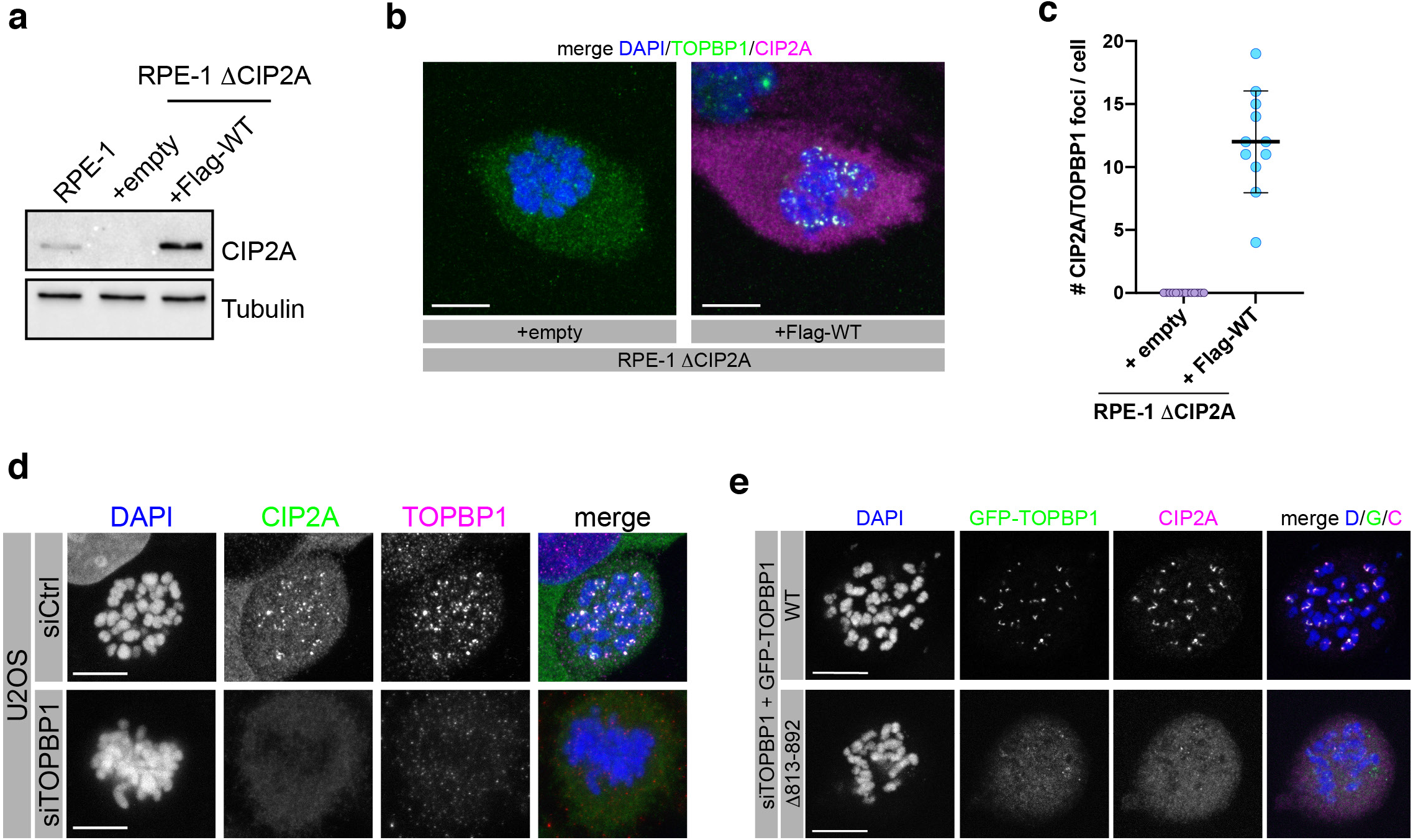
CIP2A mediates TOPBP1 DSB recruitment in mitosis. **a** Western blots of total cell extract of parental RPE-1 cells, ΔCIP2A RPE-1 cells and stably transduced ΔCIP2A cells (empty vector and vector containing Flag-tagged wild type CIP2A cDNA). **b** Confocal micrographs (maximum intensity projections) of Nocodazole-arrested empty vector (+Empty) and Flag-tagged CIP2A wild type (+Flag-WT) complemented RPE-1 ΔCIP2A cells, treated with 1 Gy of IR and stained for TOPBP1 and CIP2A. **c** Quantification of the experiment in **b**. Number of CIP2A/TOPBP1 foci per cell was assessed (n=10). **d** Confocal micrographs (maximum intensity projection) of control siRNA (siCtrl) and TOPBP1 siRNA (siTOPBP1) transfected, Nocodazole-arrested U2OS cells, 1h after irradiation with 1 Gy. **e** Confocal micrographs (maximum intensity projections) of Nocodazole-arrested GFP-TOPBP1 (wild type and Δ813-892) expressing U2OS cells, treated with 1 Gy of IR. Endogenous TOPBP1 was depleted by 5’-UTR targeting TOPBP1 siRNA. All scale bars = 10 μm.

### CIP2A-TOPBP1 recruitment to sites of mitotic DSBs is mediated by MDC1

We next investigated the role of MDC1 in the recruitment of CIP2A and TOPBP1 to sites of mitotic DSBs. We previously demonstrated that in mitosis, TOPBP1 recruitment is dependent on a direct interaction between its N-terminal BRCT1 and BRCT2 domains with the two conserved phosphorylated Serine residues S168 and S196, respectively^10^ CIP2A and TOPBP1 recruitment were significantly reduced, but not completely abrogated, in RPE-1 MDC1 knock-out cells (RPE-1 ΔMDC1) and in RPE-1 H2AX^S139A/S139A^ knock-in cells, as well as in U2OS MDC1 knock-out cells (U2OS ΔMDC1), indicating that the majority of IR-induced CIP2A/TOPBP1 structures on mitotic chromosomes are dependent on a gH2AX-MDC1 mediated recruitment mechanism (Fig. 4a,b and Supplementary Fig. 3a-c). Consistent with the notion that CIP2A acts downstream of gH2AX and MDC1 in the mitotic response to DSBs, MDC1 recruitment is not affected in RPE-1 ΔCIP2A cells (Supplementary Fig. 3d). To test if CIP2A exists in a complex with MDC1 and TOPBP1, we transfected 293FT cells with GFP-tagged MDC1, followed by immunoprecipitation with anti-GFP affinity beads.

**Fig. 4.**
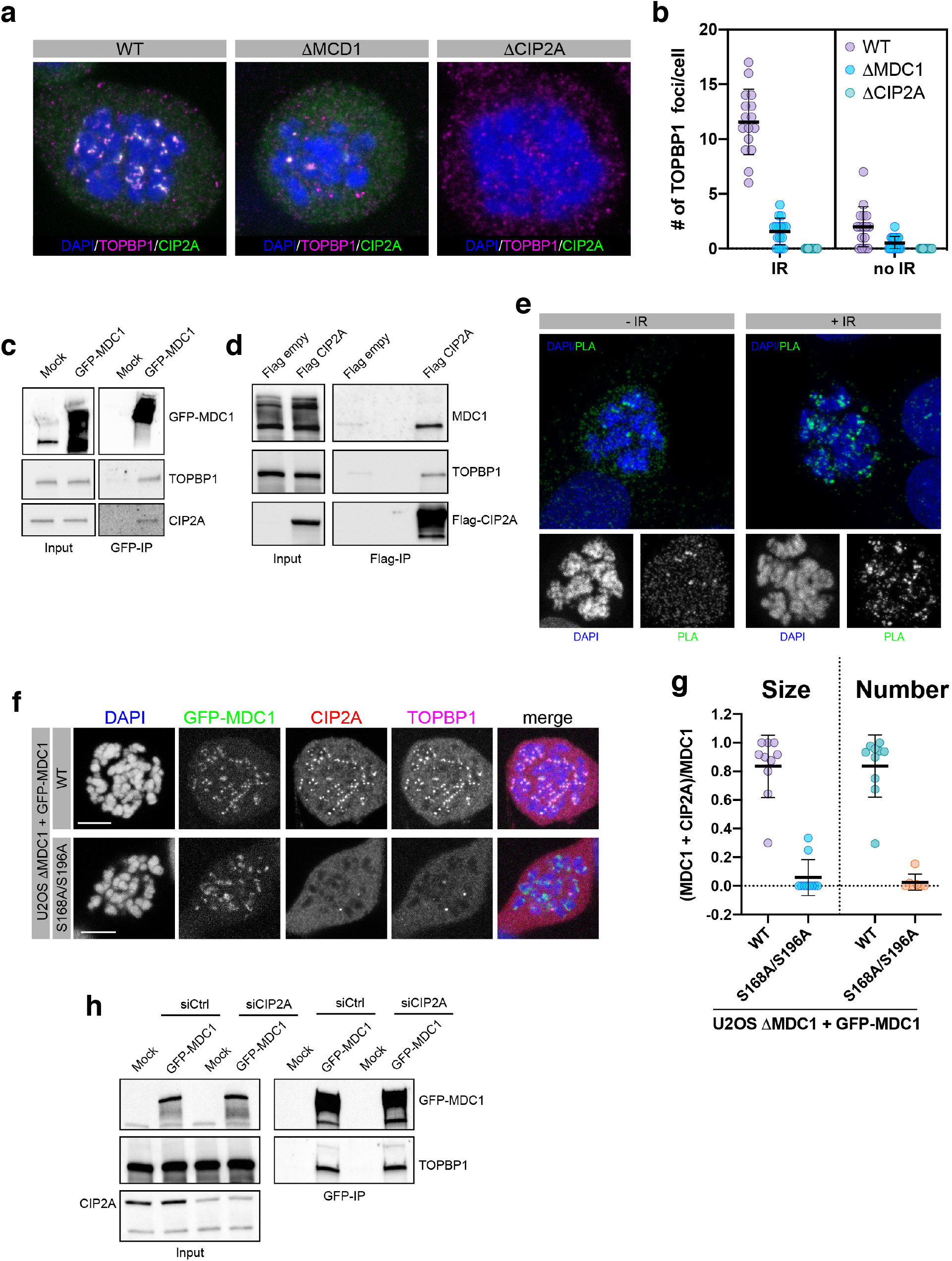
CIP2A-TOPBP recruitment to sites of mitotic DSBs is mediated by MDC1 **a** Confocal micrographs (maximum intensity projections) of Nocodazole-arrested RPE1 wild type, MDC1 knock-out (ΔMDC1) and CIP2A knock-out (ΔCIP2A) cells, irradiated with 1 Gy and stained for TOPBP1 and CIP2A. **b** Quantification of the experiment in **a**. CIP2A/TOPBP1 foci were manually counted. Each data point represents one mitotic cell n=16 per cell line and condition **c** GFP pull-downs from 293FT cells transfected with GFP-tagged full-length MDC1. **d** Flag-immunoprecipitation from 293FT cells transfected with Flag-tagged full-length CIP2A. **e** Detection of CIP2A-MDC1 co-localization by in situ PLA in U2OS cells arrested in mitosis by Nocodazole and mock treated or treated with 1 Gy of IR. **f** Confocal micrographs (maximum intensity projections) of Nocodazole-arrested U2OS ΔMDC1 cells stably transfected with GFP-tagged wild type and S168A/S196A mutated MDC1. **g** Quantitative analysis of GFP-MDC1 and CIP2A co-localization by SQUASSH: Left graph: object size colocalization (area of object overlap divided by total object area). Right graph: object number colocalization (fraction of objects in each channel that overlap ≥ 50%). Data points represent individual mitotic cells (n=9). Bars and error bars represent mean and SD. h GFP pull-down from 293FT cells transfected with GFP-tagged full-length MDC1 and either control siRNA (siCtrl) or siRNA against CIP2A (siCIP2A). All scale bars = 10 μm

Both CIP2A and TOPBP1 co-immunoprecipitated with GFP-MDC1 (Fig. 4c). Similarly, both TOPBP1 and MDC1 co-immunoprecipitated with overexpressed Flag-tagged CIP2A, thus suggesting that CIP2A and TOPBP1 exist in a ternary complex with MDC1 *in vitro* (Fig. 4d). To confirm that CIP2A is in close proximity to MDC1 also *in vivo* we performed *in situ* PLA with antibodies against CIP2A and MDC1. Similar to the results of the PLA assays with antibodies against CIP2A and TOPBP1, a clear PLA signal was detectable in mitotic cells and this signal was enriched in foci upon irradiation (Fig. 4e). Importantly, we also observed that both CIP2A and TOPBP1 foci were absent from irradiated mitotic chromosomes in U2OS ΔMDC1 cells stably expressing GFP-tagged S168A/S196A MDC1, while they were readily detectable and co-localized with MDC1 in wild type GFP-MDC1 expressing cells (Fig. 4f,g). This indicates that accumulation of both CIP2A and TOPBP1 at sites of mitotic DSBs is dependent on the direct interaction between TOPBP1 BRCT1 and BRCT2 with phosphorylated MDC1. Notably, in CIP2A depleted cells, TOPBP1 interacted as efficiently with MDC1 as in CIP2A expressing control cells, indicating that CIP2A does not simply promote the association of TOPBP1 with MDC1 (Fig. 4h). In summary, our data suggest that TOPBP1 must exist in a complex with CIP2A in order to be efficiently recruited to sites of mitotic DSBs by gH2AX-MDC1.

### CRM1-dependent nuclear export sequesters CIP2A from TOPBP1 in interphase cells

CIP2A is a 905 amino acid protein, roughly composed of two structurally distinct regions: a N-terminal Armadillo repeat domain (Arm; amino acids 1-560) and a C-terminal predicted coiled-coil region (Fig. 5a)^21^. In a proteomic screen, CIP2A was identified as a target of the chromosome region maintenance 1 (CRM1, also called exportin 1) transport receptor for the export of proteins from the nucleus^22^. Indeed, we observed that treatment of cells with two selective CRM1 inhibitors (Leptomycin B and Selinexor) led to a significant increase in nuclear localization of CIP2A in interphase cells (Fig. 5b,c). CRM1 binds to its targets via a flexible recognition motif called nuclear export signal (NES). Three NES were predicted in CIP2A based on the Eukaryotic Linear Motive Resource for Function Sites in Proteins (ELM^23^). Two of them reside in the N-terminal Arm region that mostly localized to the nucleus, thus ruling them out as genuine NES (data not shown). The third one (amino acids 598-612) is localized between the Arm repeat domain and the predicted C-terminal coiled coil region (Fig. 5a), and deletion of a fragment comprising this sequence motif (amino acids 561-625) significantly increased nuclear CIP2A concentration in interphase cells, indicating that CIP2A is a direct CRM1 ligand (Fig. 5d-f). Since TOPBP1 is predominantly nuclear, the active transportation of CIP2A from the nucleus to the cytoplasm suggested that these two proteins are spatially separated during interphase. Staining of CIP2A in cells expressing endogenously tagged Lamin A (LMNA, a component of the nuclear envelope) showed that CIP2A is not present on chromatin until nuclear export ceases upon nuclear envelope breakdown at the onset of mitosis (Fig. 5g). Consistent with the idea that CIP2A is sequestered from TOPBP1 in interphase cells, we only observed PLA signals with CIP2A and TOPBP1 antibodies in mitotic cells (Fig. 5h). Together, these data show that CIP2A and TOPBP1 are spatially separated in interphase cells by CRM1-mediated nuclear export of CIP2A.

**Fig. 5.**
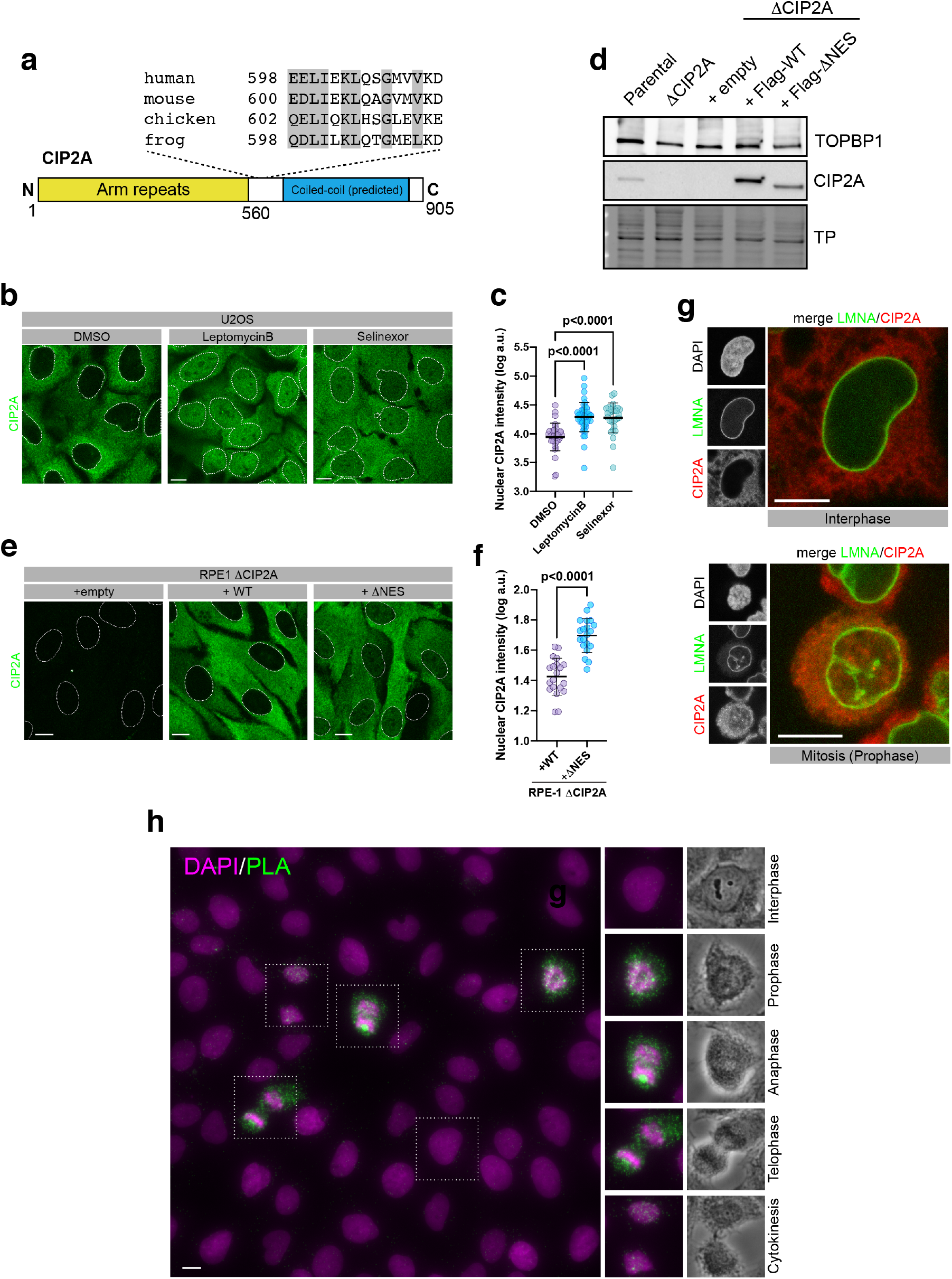
CRM1-dependent nuclear export sequesters CIP2A from TOPBP1 in interphase cells. **a** Schematic showing the structural domains of CIP2A and the location of a putative NES. Conserved amino acids matching the NES consensus motif are highlighted. **b** Confocal micrographs (maximum intensity projections) of Leptomycin B, Selinexor and control DMSO treated U2OS cells, stained for CIP2A. **c** Quantification of nuclear CIP2A staining of the experiment in **a**. Statistical significance was calculated using one-way analysis of variance and Sidak’s multiple comparison test. Bars and error bars represent mean and SD; DMSO: n=35; Leptomycin B: n=45; Selinexor: n=30. **d** Western blots of total cell extract of parental RPE-1 cells, ΔCIP2A RPE-1 cells and stably transduced ΔCIP2A cells (empty vector, Flag-tagged wild type CIP2A and Flag-tagged CIP2A ΔNES). TP stands for total protein on blot and serves as loading control. **e** Confocal micrographs (maximum intensity projections) of RPE1 ΔCIP2A cells and RPE1 ΔCIP2A cells stably transduced with Flag-tagged full-length CIP2A and CIP2A deletion mutant (human CIP2A amino acids 561-625; ΔNES), stained for CIP2A. **f** Quantification of nuclear CIP2A staining of the experiment in **e**. Statistical significance was calculated using unpaired t-test; bars and error bars represent mean and SD; n=20. **g** Confocal micrograph (maximum intensity projection) of Hela cells expressing endogenous Clover-tagged LMNA and stained for CIP2A. **h** Micrograph of *in situ* PLA, using antibodies against CIP2A and TOPBP1, in unsynchronized and untreated U2OS cells. DAPI staining and phase contrast were used to visualize interphase and four mitotic stages. All scale bars = 10 μm.

### CIP2A-independent TOPBP1 foci formation in interphase cells

An important prediction from the spatial separation of CIP2A from TOPBP1 in interphase cells is that CIP2A is not implicated in TOPBP1 recruitment to sites of DSBs in interphase. Indeed, in response to DSB induction by IR, TOPBP1 foci formation in CIP2A deficient interphase RPE-1 and U2OS cells is indistinguishable from cells that express CIP2A (Fig. 6a,b and Supplementary Fig. 4a,b). Moreover, CIP2A is also not implicated in Treacle-mediated TOPBP1 accumulation in nucleolar caps after targeted induction of rDNA breakage (Fig. 6c,d). It is currently unclear if CIP2A interferes with DSB responses in interphase cells and thus needs to be sequestered from chromatin by nuclear export. Its presence in the nucleus of interphase cells does not seem to negatively impact on TOPBP1 recruitment to sites of DSBs (Supplementary Fig. 4c). Interestingly, we observed recruitment of CIP2A to foci and co-localization with TOPBP1 in a subset of Leptomycin B or Selinexor treated interphase cells (Fig. 6e,f). This suggests that CIP2A, when forced in the nucleus in interphase cells, is capable to bind to TOPBP1 at sites of DSBs, arguing against the possibility that CIP2A-TOPBP1 association is directly regulated during the cell cycle, but rather suggests that the regulation occurs via spatial separation in interphase due to nuclear export.

**Fig. 6.**
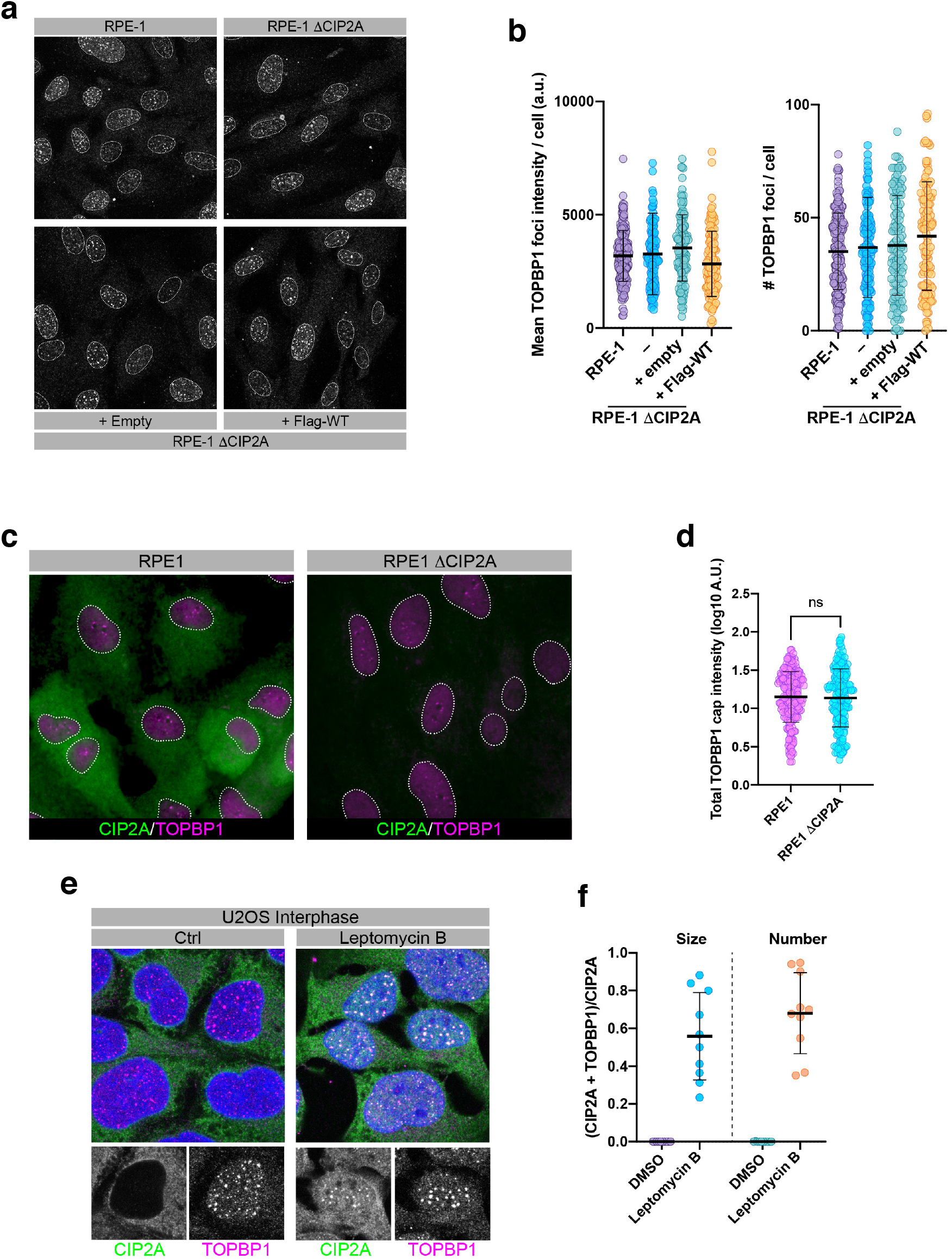
CIP2A-independent TOPBP1 foci formation in interphase cells. **a** Confocal micrographs of unsynchronized parental RPE-1 cells, RPE-1 ΔCIP2A cells and RPE-1 ΔCIP2A cells stably transduced with empty vector (+Empty) or Flag-tagged full-length CIP2A (+Flag-WT), treated with 3 Gy of IR and stained for TOPBP1. **b** Quantification of TOPBP1 foci mean intensity and number of TOPBP1 foci per nucleus in interphase parental RPE1 cells, RPE1 ΔCIP2A cells and RPE1 ΔCIP2A cells stably transduced with empty vector (+Empty) or Flag-tagged full-length CIP2A (+Flag-WT). Bars represent mean and SD; RPE-1: n=200; ΔCIP2A: n=144; +Empty: n=128; +Flag-WT: n=125. **c** Micrographs of RPE1 wild type and ΔCIP2A cells transfected with I-Ppo1 mRNA to induce rDNA breaks and stained for CIP2A and TOPBP1. **d** Quantification of the experiment in **c**. Statistical significance was calculated using unpaired t-test. Bars and error bars represent mean and SD; n=323 (RPE1), 329 (RPE1 ΔCIP2A). ns = not significant. **e** Confocal micrographs (maximum intensity projections) of U2OS cells pre-treated with DMSO and Leptomycin B, irradiated with 3 Gy and stained for CIP2A and TOPBP1. **f** Quantitative analysis of CIP2A and TOPBP1 colocalization of the experiment in **e** by SQUASSH. Left: object size colocalization (area of object overlap divided by total object area). Right: object number colocalization (fraction of objects in each channel that overlap ≥ 50%). Each data point represents one cell (n=10). Bars and error bars represent mean and SD. All scale bars = 10 μm.

### CIP2A deficient cells display increased micronuclei formation, chromosomal instability and DSB repair defects

Cells unable to recruit TOPBP1 in mitosis due to disrupted MDC1-TOPBP1 interaction show increased micronuclei (MNi) formation and chromosomal instability^10^, most likely due to the inability to properly segregate acentric chromosome fragments^24^. In line with its capacity to control TOPBP1 localization in mitosis, CIP2A deficient cells also displayed increased spontaneous MNi formation, which was slightly increased after IR treatment, and was rescued by re-expression of wild type CIP2A (Fig. 7a). The majority of MNi stained negative for CENPA, a centromeric marker, indicating that they mostly contain acentric chromosome fragments (Fig. 7b). Consequently, we also observed a significantly increased number of abnormal structures in metaphase spreads of RPE-1 ΔCIP2A cells, but not in RPE-1 ΔCIP2A cells stably transduced with wild type CIP2A (Fig. 7c,d). To explore whether loss of CIP2A affects DSB repair in the subsequent interphase after irradiation in mitosis, we first synchronized parental RPE-1 cells and ΔCIP2A cells as well as ΔCIP2A cells stably expressing Flag-tagged wild type CIP2A in mitosis followed by irradiation and release them from the mitotic block. gH2AX foci (a marker for unrepaired DSBs) were quantified 24 h post-irradiation. There was a significant increase in residual gH2AX foci in cells deficient for CIP2A 24 h post-irradiation and re-expression of wild type CIP2A rescued this phenotype (Fig. 7e,f). We conclude from these results that CIP2A controls TOPBP1 recruitment to sites of mitotic DSBs and that in its absence, cells accumulate MNi and chromosomal instability.

**Fig. 7.**
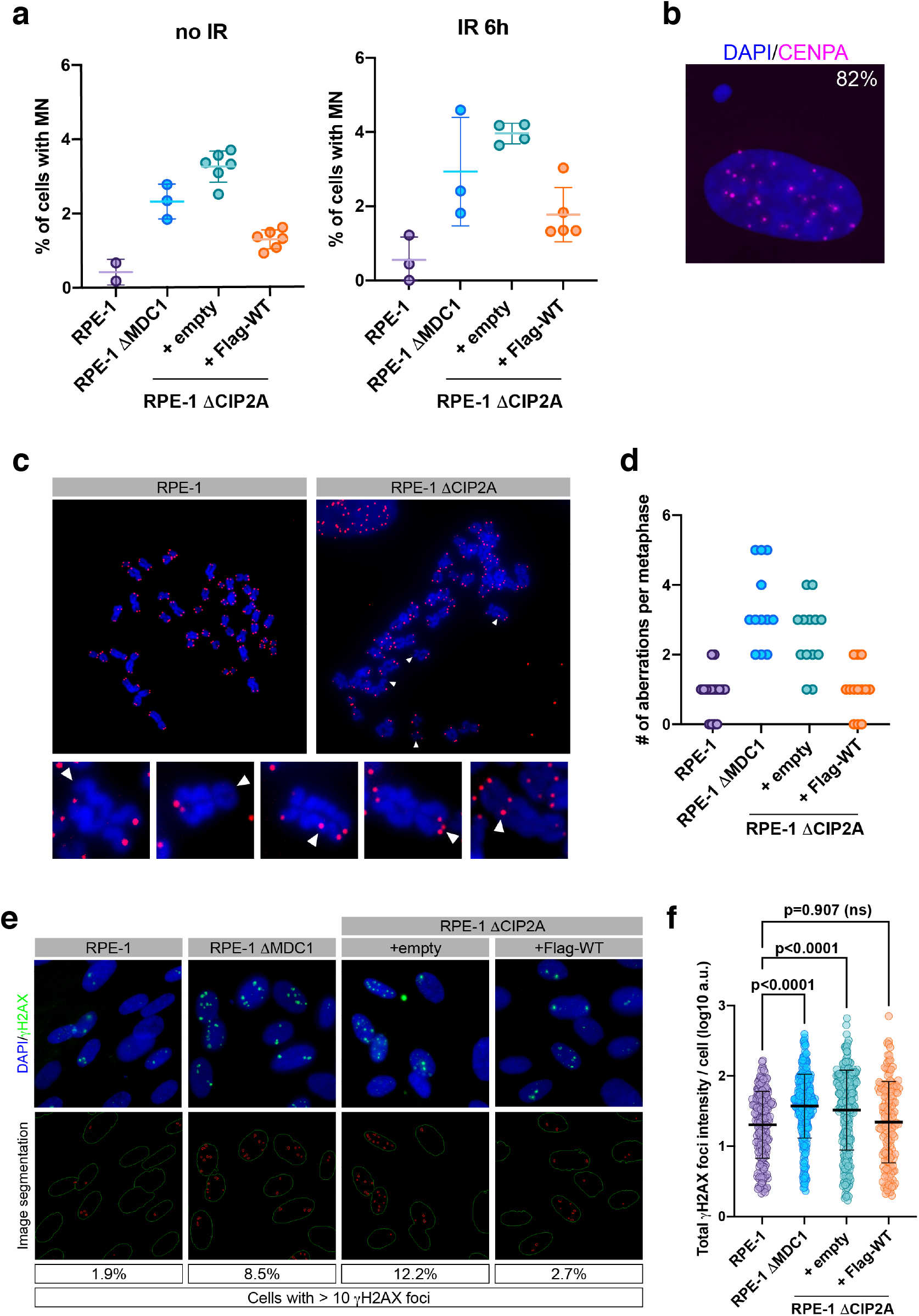
CIP2A deficient cells display increased micronuclei formation, chromosomal instability and DSB repair defects. **a** Quantification of micronuclei formation in non-irradiated and irradiated RPE-1 parental cells, RPE-1 ΔMDC1 cells, RPE-1 ΔCIP2A cells stably transduced with empty vector (+ empty) and Flag-tagged wild type CIP2A (+ Flag-WT). **b** Example of RPE1 ΔCIP2A cell with micronucleus, stained with CENPA antibody. Percentage of CENPA-negative micronuclei are indicated. **c** Examples of chromosomal aberrations in metaphase spreads derived from RPE-1 and RPE-1 ΔCIP2A cells. Aberrations include single chromatid telomere loss, sister chromatid telomere loss, interstitial telomeres, telomere duplications and dicentric chromosomes. **d** Quantification of chromosomal aberrations in metaphase spreads derived from untreated RPE-1 parental cells, RPE-1 ΔMDC1 cells and RPE-1 ΔCIP2A cells stably transduced with empty vector (+ empty) and Flag-tagged wild type CIP2A (+ Flag-WT). Aberrations were counted manually. Each data point represents one metaphase. n=16 (WT), 12 (ΔMDC1), 13 (+Empty), 13 (+Flag-WT) **e** Residual gH2AX foci 24 h after irradiation of RPE-1 parental cells, RPE-1 ΔMDC1 cells and RPE-1 ΔCIP2A cells stably transduced with empty vector (+ empty) and Flag-tagged wild type CIP2A (+ Flag-WT). Cells were arrested in G2 by RO-3306, irradiated with 0.5 Gy and released from the mitotic arrest. Upper panels: representative micrographs of cells stained for gH2AX. Lower panels: results of image segmentation for quantification. Percentage of cells with > 10 gH2AX foci are indicated. **f** Quantification of the experiment in **e**. Statistical significance was calculated using one-way analysis of variance and Sidak’s multiple comparison test. Bars and error bars represent mean and SD; RPE1: n=368; ΔMDC1: n=330; +Empty: n=219; +Flag-WT: n=224. All scale bars=10 μm.

## CONCLUSION

In summary, our findings demonstrate that CIP2A functions downstream of MDC1 to mediate TOPBP1 recruitment to sites of chromosome breaks in mitosis, which is critical for proper segregation of acentric chromosome fragments. As CIP2A deficiency gives rise to DSB repair defects after irradiation of mitotic cells, it is possible that CIP2A mediates productive DSB repair by stabilizing and/or tethering broken chromosomes during cell division until they enter the subsequent cell cycle where these lesions can be repaired. We have also shown that the effect of CIP2A on TOPBP1 is restricted to mitosis by nuclear exclusion, which is similar to the spatial control of the GEN1 Holliday junction resolvase whose activity is needed at the onset of mitosis but causes elevated crossover formation when present in the nucleus in interphase^25^. Cancer cells frequently have to deal with remnants of replication problems from the preceding S-phase, either as a consequence of oncogene-driven cell cycles, or as a result of defective HR pathways due to the loss of tumor suppressor genes such as BRCA1 and BRCA2. We surmise that at least a subset of these replication intermediates will be converted to DSBs in mitosis and will thus require the CIP2A-TOPBP1 complex for their stabilization. Indeed, in a complementary work submitted elsewhere, we found that loss of CIP2A or disruption of the CIP2A-TOPBP1 complex is lethal in HR-deficient cells, most likely due to rampant mis-segregation of acentric chromatin fragments that occur as a result of aberrant DNA replication in these cells (Adam et al., *submitted).* Collectively, our data firmly establish CIP2A-TOPBP1 as a physiologically critical component of the cellular machinery that deals with DNA lesions specifically during mitosis.

## METHODS

### Cell culture

293T, U2OS, RPE-1 hTERT and HeLa cells were cultured in Dulbecco’s modified Eagle’s medium (DMEM Gibco™, 61965026), supplemented with 10% fetal calf serum (PAN Biotech, P40-37500), 2 mM L-glutamine and penicillin-streptomycin antibiotics under standard cell culture conditions in a CO2 incubator (37 °C; 5 % CO2). Stably transfected U2OS cell lines were cultured in the presence of 400 μg/mL Geneticin (G418 sulfate, Sigma, G-3632). RPE-1 ΔCIP2A cell line complemented with lentiviral plasmids were cultured in the presence of 50 μg/mL Nourseothricin dihydrogen sulfate (Roth, 3011). The generation of U2OS and RPE1 ΔMDC1 knock-out cell lines and complemented derivatives as well as RPE1 H2AX^S139A/S139A^ knock-in cell lines were previously described^10 26^. Irradiation of cells was done in a YXLON, Y.SMART583 custom-made X-ray device. All cell lines were regularly tested for mycoplasma contamination.

### Generation of a CIP2A knock-out RPE-1 cell line

RPE1 hTERT cells were cultured overnight at a density of 0.1 x 10^6^ with 1 mL DMEM growth medium (Gibco™, 61965026), supplemented with 10 % FBS (PAN Biotech, P40-37500), and Antibiotic-Antimycotic (Gibco™, 15240062). For transfection, 375 ng of synthetic sgRNA (TrueGuide™, CRISPR932782_SGM) along with 6 μL Cas9 mRNA (Invitrogen™, A29378), and 18 μL of the MessengerMAX™ reagent (Invitrogen™, LMRNA015) were separately diluted in 300 μL Opti-MEM™ I (Gibco™, 31985047) each. After 10 min of incubation at room temperature, both mixtures were combined and added to the plate at 100 μL per well. The cells were kept in culture for another 48 h, trypsinized and transferred to a 96-well plate in serial two-fold dilution, starting with 4’000 counts. Single clones were successively identified by eye under a transluminescence microscope and progressively expanded to larger culture volumes. One successful knock-out could be identified by means of immunofluorescence as well as western blotting total cell extracts. An Adenine insertion at position 5110 of the gene was confirmed by subcloning the targeted area into a pCR™4Blunt-TOPO^®^ vector (Invitrogen™, 450031) and subsequent colony sequencing of transformed Stellar™ Competent Cells (Takara, 636766). gRNA sequence: GGTCTTCAAGTAGCTCTACA

### Generation of endogenous-tagged Clover-LMNA Knock-In cell lines

The nuclear lamina protein LMNA was endogenously labeled, in both HeLa and U2OS cell lines, with the monomeric green fluorescent protein Clover as previously described^27^. Briefly, cells were co-transfected with a Clover donor plasmid, that incorporates sequence homology to the locus adjacent to LMNA-start on each side of the tag, and one that encodes for Cas9 along with a chimeric gRNA, targeting two sites within the same region The cultures were expanded to confluency, detached using trypsin-EDTA (Gibco, 15400054), washed, and resuspended in 1.5 mL PBS buffer (Gibco, 10010015), supplemented with 1 % BSA (Sigma-Aldrich, A7906), 25 mM HEPES (Gibco, 15630056), and 1 mM EDTA (AppliChem, A4892). The suspension was then passed through the cell strainer cap of a 5 mL PS test tube (Falcon, 352235). Single cells expressing relevant amounts of green fluorescence were isolated and collected in a cell sorter (BD Biosciences, FACSAria III 4L).

### Cloning and Mutagenesis

pIRESneo2-EGFP-TOPBP1 plasmids are described in^12^. pIRESneo2-EGFP-TOPBP1-ΔBRCT6 was generated by site directed mutagenesis of pIRESneo2-EGFP-TOPBP1-WT using QuikChange II XL Site-Directed Mutagenesis kit (Agilent Technologies, 200522).

Sequences of the mutagenesis primers for TOPBP1-ΔBRCT6 were, forward: 5’-CCC AAA ATG AGC TTG GAT ATC AGC GCA-3’; reverse:

5’ - TTC CTT CTC TGA CTG GGC CTC TTT CAG-3’. Generation of pIRESneo2-EFGP-TOPBP1-W1145R is described elsewhere^14^ pIRESneo2-EGFP-TOPBP1-Δ813-892 was generated by cloning a fragment of TOPBP1 lacking aa 813 to aa 892 that was generated synthetically by Genscript (genscript.com) into pIRESneo2-EGFP-TOPBP1-WT by using SbfI and StuI restriction enzymes. pHIV-NAT-T2A plasmids containing Flag-tagged fulllength CIP2A cDNA are described in Adam et al. (submitted). Deletion of the residues 561-625 (ΔNES) was done by deletion PCR using Phusion^®^ High-Fidelity DNA Polymerase (NEB), followed by DpnI digestion of the template and transformation into bacteria. Primers were designed using the Quick Change primer design tool (Agilent) and here are the sequences: forward: 5’-AAA CAA TGC CTA TAG ACA ACA GGA ATA TGA AAT GAA ACT ATC CAC ATT AG-3’; reverse:

5’-CTA ATG TGG ATA GTT TCA TTT CAT ATT CCT GTT GTC TAT AGG CAT TGT TT-3’.

Plasmids pIRES V5 I-Ppo1 for I-Ppo1 mRNA production and purification were described^28^

### Lentivirus production

293T cells were plated at 80 % confluence in a 10 cm dish with 8 mL of culture medium one day prior to transfection. Transfection was carried out using Lenti-X Packaging Single Shot (VSV-G) system (Takara Bio, 631275) according to the following protocol: 7 μg of lentiviral plasmid were diluted in 600 μL of sterile water, incubated at room temperature for 10 min and added to the cells dropwise. 293T infected cells were supplemented with fresh culture medium 4 h after infection and further incubated for 48 and 72 h before proceeding with the first and second lentivirus harvest, respectively. Lentivirus-containing supernatants were centrifuged at 4 °C for 10 min at 500 g and passed through a 0.45 μm filter. Virus titer was measured using the QuickTiter™ Lentivirus Titer Kit (Cell Biolabs, VPK-107-T) according to the manufacturer’s instructions.

### Transduction of RPE-1 ΔCIP2A cells

RPE-1 ΔCIP2A cells were plated in 6 cm dishes at 80 % confluence in culture medium one day prior to infection. Cells were incubated with virus-containing supernatants in the presence of 8 μg/mL Polybrene (Santa Cruz Biotech, sc-134220) for 24 h. The virus volume needed for infection was calculated using the following formula: Total Transduction Units (TU) needed = cells seeded*MOI (in this experiment MOI = 5 was used).

Total mL of lentiviral particles to add to each 6 cm dish = (Total TU needed) / (TU/mL) After incubation, virus-containing medium was discarded and replaced with fresh culture medium. Cells were incubated for an additional 48 h before proceeding with protein analysis to evaluate expression of the gene of interest. Transduced cells were selected in the presence of 100 μg/mL Nourseothricin dihydrogen sulfate.

### Cell transfection with RNA

The control siRNA (siCtrl), siRNA targeting TOPBP1 (siTOPBP1) and the 3’ UTR sequence of TOPBP1 gene (siTOPBP1 3’UTR) were obtained from Microsynth AG. The sequences of the siRNAs are the following:

siCtrl: UGGUUUACAUGUCGACUAA-dTdT,

siTOPBP1: ACAAAUACAUGGCUGGUUA-dTdT

siTOPBP1 3’UTR: GUAAAUAUCUGAAGCUGUATT-dTdT

ON-Target plus SMART pool siRNA targeting CIP2A was purchased from Dharmacon (L-014135-01-0005) and contains a mix of four different siRNAs:

Sequence 1: ACAGAACUCACACGACUA

Sequence 2: GUCUAGGAUUAUUGGCAAA

Sequence 3: GAACAAAGGUUGCAGAUUC

Sequence 4: GCAGAGUGAUAUUGAGCAU For siRNA transfection, cells were grown in a six-well plate 24 h prior to transfection. 60 % confluent cells were transfected using 20 nM siRNA and Lipofectamine RNAiMAX (Invitrogen, 13778150) according to the manufacturer’s instructions. Cells were incubated with siRNA-lipid complexes for 48 h before being seeded on coverslips and harvested 72 h after transfection. I-Ppo1 mRNA transfection was done using the Lipofectamine MessengerMax Reagent (Invitrogen™, LMRNA015) according to the manufacturer’s protocol.

### SDS-PAGE and western blotting

SDS-PAGE and western blotting were performed using 4-20 % Mini-PROTEAN TGX Stain-free Precast Gels (Bio-Rad, 4568094) and Trans-Blot Turbo 0.2 μm nitrocellulose membranes (Bio-Rad. 1704158), respectively. Antibodies against the following antigens were used at the indicated dilutions: GFP (Mouse, 11814460001, Roche, 1/5000), HA (Rabbit, ab9110, Abcam, 1/4000), MDC1 (Rabbit, ab11171, Abcam, 1/5000), TOPBP1 (Rabbit, ab2402, Abcam, 1/1500) and CIP2A (Rabbit, 14805, Cell Signaling, 1/1000). Blots were developed with SuperSignal™ West Femto Maximum Sensitivity Substrate (Thermo Scientific™, 34096) and image acquisition was carried out using ChemiDoc MP Imaging System (Bio-Rad, 17001402).

### HA Immunoprecipitation

For preparation of lysates for immunoprecipitations (IPs), cells were washed in cold PBS buffer pH 7.45 (Gibco™, 18912-014), harvested with a cell scraper and centrifuged at 500 g for 3 min. PBS was then discarded and cells were resuspended in an appropriate volume of IP lysis buffer (50mM Tris-HCl pH 7.6, 150 mM NaCl, 5 mM MgCl_2_, 0.5 % Triton X-100), supplemented with cOmplete EDTA-free protease inhibitor cocktail (Roche, 11873580001) and 25 U/ml Benzonase nuclease (Sigma, E1014). Cells were incubated on ice for 30 min and lysates were cleared by centrifugation at 16,000 g for 15 min. 2 mg of the soluble protein fraction were then transferred into fresh tubes and incubated with 30 μL of monoclonal antiHA agarose beads (Sigma, A2095). Samples were incubated for 2 h with end-over-end rotation at 4 °C. Immunoglobulin-antigen complexes were washed 3x 15 min in cold PBS with end-over-end rotation before elution in 2X SDS sample buffer (Geneaid, PLD001, 100 mM DTT).

### GFP Immunoprecipitation

Preparation of lysates was performed as described for HA-immunoprecipitation. 1 mg of lysate was incubated with 25 μL of equilibrated GFP-trap magnetic agarose beads (Chromotek, gtma-20) for 1 h with end-over-end rotation at 4 °C. Samples were washed 4 x 10 min in cold PBS with end-over-end rotation before elution in 2X SDS sample buffer.

### FLAG Immunoprecipitation

For preparation of lysates, cells were washed in cold PBS buffer pH 7.45, scraped and centrifuged at 500 g for 3 min. PBS was then discarded and cells were resuspended in an appropriate volume of IP lysis buffer (50 mM Tris-HCl pH 7.4, 150 mM NaCl, 1 mM EDTA, 1 % Tryton X-100) supplemented with cOmplete EDTA-free protease inhibitor cocktail and 25 U/mL Benzonase nuclease. Lysates were cleared by centrifugation and 400 μg of protein lysate were incubated with 40 μL of Anti-FLAG^®^ M2 affinity gel (Sigma, A2220) for 2 h with end-over-end rotation at 4 °C. Immunoglobulin antigen complexes were washed 3x 15 min in cold TBS with end-over-end rotation before elution in 2X SDS sample buffer.

### On-beads digestion and Mass Spectrometry

The Functional Genomics Center of the University of Zurich (FGCZ) was commissioned to perform the mass spectrometry analysis. HA immunoprecipitation was performed as described above. After the last wash, precipitated material was resuspended in PBS buffer pH 7.45 and subjected to on-beads trypsin digestion according to the following protocol. Beads were washed twice with digestion buffer (10 mM Tris, 2 mM CaCl_2_, pH 8.2). After the last wash, the buffer was discarded and monoclonal anti-HA agarose beads were resuspended in 10 mM Tris, 2 mM CaCl_2_, pH 8.2 buffer supplemented with trypsin (100 ng/μL in 10 mM HCl). The pH was adjusted to 8.2 by the addition of 1 M Tris pH 8.2. Digestion was performed at 60 °C for 30 min. Afterwards, supernatants were collected, and peptides were extracted from beads using 150 μL of 0.1 % trifluoroacetic acid (TFA). Digested samples were dried and reconstituted in 20 μL ddH2O + 0.1 % formic acid before performing Liquid chromatography-mass spectrometry analysis (LCMS/MS). For the analysis 1 μL were injected on a nanoACQUITY UPLC system coupled to a Q-Exactive mass spectrometer (Thermo). MS data were processed for identification using the Mascot search engine (Matrixscience) and the spectra was searched against the Swissprot protein database (human and mice, retrieved 18/09/2020).

### Immunofluorescence

Cells were grown on glass coverslips and washed two times with cold PBS before fixation with cold methanol for 10 min on ice. Methanol was discarded and cells were washed two times with PBS at room temperature and incubated with blocking buffer (10% FCS in PBS) for at least 1 hour. Primary antibody incubations were performed at 4 °C overnight. Coverslips were then washed 3x 10 min with PBS and secondary antibody incubation was performed for 1 h at room temperature in the dark. After washing 3x 10 min with PBS, coverslips were mounted on glass microscopy slides (Thermo Scientific, 630-1985, dimensions L76 X W26 mm) with VECTASHIELD^®^ mounting medium containing 0.5 μg/mL 4’,6-diamidino-2-phenylindole dihydrochloride (DAPI) (Vector Laboratories, H-1200). The following antibodies were used at the indicated dilutions: CIP2A (Mouse, sc-80659, Santa Cruz, 1/800), Cyclin A (Mouse, 611269, BD Biosciences, 1/100), γH2AX (Mouse, 05-636-I, Merck, 1/500), MDC1 (Rabbit, ab11171, Abcam, 1/300), TOPBP1 (Rabbit, ABE1463, Millipore, 1/300) CENPA (Mouse, ab1339, Abcam, 1/500).

### Widefield microscopy

Widefield image acquisition was done on a Leica DMI6000B inverted fluorescence microscope, equipped with Leica K5 sCMOS fluorescence camera (16-bit, 2048 x 2048 pixel, 4.2 MP) and Las X software version 3.7.2.22383. A HC Plan Apochromat 40X/0.95 PH dry objective and HCX Plan Apochromat 63X/1.40, and Plan Apochromat 100X/1.40 PH oil immersion objectives were used for image acquisition. For triple-wavelength emission detection, we combined DAPI with EGFP or Alexa Fluor 488 and Alexa Fluor 568.

### Confocal microscopy

Confocal images were acquired with a Leica SP8 confocal laser scanning microscope coupled to a Leica DMI6000B inverted stand, with a 63x, 1.4-NA Plan-Apochromat oil-immersion objective. For quadruple-wavelength emission detection we combined DAPI with EGFP, Alexa Fluor 568 and Alexa Fluor 647. For triple-wavelength emission detection we combined DAPI with EGFP or Alexa Fluor 488, and Alexa Fluor 568. The sequential scanning mode was applied, and the number of overexposed pixels was kept at a minimum. 7 to10 z-sections were recorded with optimal distances based on Nyquist criterion. For optimal representation in figures, maximum intensity projections were calculated using Fiji^29^. For quantitative assessment of protein colocalization, the SQUASSH plugin (part of the MosaicSuite) for ImageJ and Fiji was used^10,30^

### Image processing

Unprocessed grayscale tagged image files (TIFs) and maximum intensity projections of confocal z-stacks were exported from Fiji, followed by pseudo-coloring and adjustment of exposure or brightness/contrast in Adobe Photoshop CC2020. For maximum data transparency and preservation, unprocessed grayscale images or maximum intensity projections were imported as smart objects and adjustments were done using adjustment layers. Processed images were saved as multilayer PSD files.

### Quantification of TOPBP1 foci in interphase cells

Unsynchronized cells were grown on coverslips under standard culture conditions. Shortly before irradiation with 3 Gy, the culture medium was changed and replaced with fresh medium containing either 10 μM Leptomycin B (Apollo Scientific, BIL2101) or 10 μM Selinexor (Focus Biomolecules, 10-4011-0005). For control cells, the medium was replaced with fresh medium containing DMSO. Irradiated cells were then further incubated for 3 h to achieve inhibition of CRM-1 mediated nuclear export of CIP2A and recruitment of TOPBP1 to DNA damage foci. Cells were then washed twice in cold PBS before fixation with ice-cold methanol and staining with anti-TOPBP1 and anti-CIP2A antibodies. Image acquisition was done using the Leica SP8 confocal microscope. Quantification was done using CellProfiler 3.0^31^ (CellProfiler analysis pipeline available upon request). The data was processed with R (r-project.org) and graphs were generated with GraphPad Prism 9.0

### Quantification of γH2AX foci

For quantification of γH2AX foci, cells were grown on coverslips and incubated with RO-3306 (Sigma, SML0569) for 16 h to synchronize cell population in late G2. Cells were subsequently released from the cell cycle arrest by washing 3x 5 min with warm PBS and by adding fresh culture medium. Cells were further incubated for 20 min to allow them to progress into mitosis before irradiation with 0.5 Gy, when indicated, and then fixed 24 h after irradiation with ice-cold methanol. Cells were stained with mouse monoclonal anti-γH2AX antibody as described above. Image acquisition was done using widefield microscopy and quantification was done using CellProfiler 3.0^31^ (CellProfiler analysis pipeline available upon request). The data was processed with R (r-project.org) and graphs were generated with GraphPad Prism 9.0

### Quantification of micronuclei

For quantification of micronuclei formation after irradiation, unsynchronized cell populations were grown on coverslips and either treated with 3 Gy or left untreated. 6 h and 24 h after IR cells were washed in cold PBS and fixed with ice-cold methanol for 10 min. Methanol was discarded and cells were washed twice before mounting the coverslips on microscopy glass slides with VECTRASHIELD containing DAPI. Images were captured by widefield microscopy and more then 1000 cells were taken into consideration for quantification. Micronuclei were counted manually in Fiji, and nuclei were counted using the CellProfiler 3.0 software (CellProfiler analysis pipeline available upon request). For quantification of CENP-A positive and negative MNi, cells were stained with mouse monoclonal anti-CENP-A antibody, following standard immunofluorescence protocol, as described above. Image acquisition was done using widefield microscopy and quantification of CENP-A positive and negative MNi was done manually.

### *In Situ* PLA assay

U2OS cells were seeded on round glass coverslips (Thermo Scientific Menzel, CB00120RA120) in 24-well plates (TPP, 92024) at a density of 0.8 x 10^5^ per well. When indicated, cells were incubated with 16.6 μM nocodazole (Sigma-Aldrich, M1404) for another 15 h to arrest them in prometaphase. Half of the samples were then irradiated with a dose of 1 Gy and placed back to the incubator for 1 h. The growth medium was removed and all coverslips were washed with PBS, fixed with ice-cold methanol on ice for 12 min, and washed again 3x in PBS. All subsequent steps were performed using Duolink^®^ PLA Reagents (Sigma-Aldrich, DUO92001, DUO92005, DUO92008, DUO82049). The blocking solution was applied for 1 h in the humidified incubator at 37 °C. The following antibody combinations were used: mouse-anti-CIP2A (Santa Cruz, 2G10-35B; 1/800) and rabbit-anti-TOPBP1 (Millipore, ABE1463; 1/250) or rabbit-anti-MDC1 (Abcam, ab11171; 1/200). Antibody incubation was done overnight at 4 °C under humid conditions. PLUS and MINUS PLA probes were prepared according to the manufacturer’s protocol and applied analogous to the previous step but for 1 h at 37 °C in the humidified incubator. Cells were washed for 15 min, then 10 min with wash buffer B, followed by 5 min with 0.01X wash buffer B, and mounted on frosted glass slides using VECTASHIELD PLUS antifade mounting medium with DAPI (Adipogen, VC-H-2000).

### Airyscan high-resolution confocal microscopy

Cells were grown on high precision glass coverslips #1.5H, 0.17 mm thick (Assistent) and either synchronized in late G2 with RO-3306, as described above or in prometaphase using 100 ng/mL nocodazole for 16 h. Cells synchronized with RO-3306 were released into mitosis and irradiated with 1 Gy. Irradiated cells were further incubated at 37°C and fixed at 30, 45 and 60 min after irradiation in order to collect prophase, metaphase and anaphase cells, respectively. Nocodazole arrested cells were irradiated with 1 Gy and incubated for 1 h at 37°C before fixation. Cells were washed twice in cold PBS before fixation with ice-cold methanol for 10 min on ice. Methanol was discarded and cells were then washed twice with PBS at room temperature before incubation with blocking buffer (10% FCS in PBS) for 2 h. Primary antibodies were diluted in 5% FCS in PBS and incubated overnight at 4 °C. Cells were then washed 3x 10 min with blocking buffer before incubation with Alexa Fluor secondary antibodies (diluted 1/1000 in 5% FCS) for 1h at room temperature in the dark. Secondary antibody was then washed 3x 10 min with PBS. Cells were then stained with 1 μg/mL DAPI (AppliChem, A1001) diluted in PBS for 15 min at room temperature in the dark. DAPI was discarded and cells were further washed 2x 5 min with deionized water to completely remove the PBS. Coverslips were then mounted on microscopy glass slides without frosted edges (R. Langenbrick GmbH, Dimensions: L 76 x W 26 mm) using VECTASHIELD HardSet Antifade Mounting Medium (Vector Laboratories, H-1400). Airyscan confocal imaging was carried out at the ScopeM imaging facility of ETH Zurich, using an LSM 880 Airyscan inverted microscope (ZEISS) equipped with a DIC Plan Apochromat 63x/1.4 oil immersion objective and an Airyscan 32-pinhole detector unit. DAPI was detected using a 405-nm diode laser and 420-480 nm plus 495-550 nm band pass emission filters, Alexa 488 and GFP were detected with the 488-nm line of an argon laser and 420-480 nm plus 495-550 nm band pass emission filters, and Alexa 568 was detected using a 561-nm diode-pumped solid-state laser and 495-550 nm band pass plus 570 nm long pass emission filters. Z-stacks of entire cells were acquired using optimal step size. Raw data were processed using Airyscan processing with Wiener Filter (auto setting) available in ZEN Black software version 2.1, yielding 8-bit images with approximately 180 nm lateral resolution. Display of images was adjusted for intensity for optimal display of structures of interest.

### Metaphase analysis

Cells were grown to 90% confluence on a 10 cm plate and were treated with 0.1 μg/mL KaryoMax Colcemid (Thermo Fisher Scientific) for 16 h overnight. Cells were trypsinized and transferred to a 15 ml Falcon^™^ tube, centrifuged at 176 g for 5 min and carefully resuspended in 5 ml of pre-warmed hypotonic buffer (15 % FBS, 75 mM KCl) with intermittent agitation and incubated for 15 min at 37 °C. Cells were again pelleted at 176 g for 5 min, the supernatant was discarded, and the cell pellet was resuspended in 200 μl of hypotonic buffer. Cells were fixed by adding drop-wise 7 ml of ice-cold MeOH:AcOH 3:1 while slowly vortexing followed by 40 min on ice. After centrifugation at 176 g for 5 min, supernatant was discarded, and cells were resuspended in remaining 200 μl of the fixation buffer. 20-25 μl of the cell suspension was then dropped at a 45° angle onto a wet glass slide and air-dried for 10 min. Metaphases were stained with DAPI (VECTASHIELD containing DAPI), covered with a glass coverslip and sealed with nail varnish. Telomere FISH was conducted using the Telomere PNA FISH Kit/Cy3 (Dako, K5326) according to the protocol provided by the supplier.

## Acknowledgments

We thank Thanos Halazonetis, Steve Jackson, Brian McStay and Graham Dellaire for providing valuable reagents, and Andrew Blackford for sharing unpublished observations. Imaging was performed with equipment maintained by the Center for Microscopy and Image Analysis, University of Zürich or the Scientific Center for Optical and Electron Microscopy (ScopeM) of the ETH Zürich. Cell sorting was carried out by the Flow Cytometry Core Facilities at the University of Zürich. Mass spectrometry was carried out by the Functional Genomics Center of the University of Zürich. The Durocher lab is supported by grants from the Canadian Institutes for Health Research (FDN143343) and the Canadian Cancer Society (705644). The Stucki lab is supported by two project grants from the Swiss National Foundation (31003A_163141 and 310030_189141) and by the Kanton of Zürich.

## Author contributions

The project was conceived and supervised by M.S. and D.D. Biochemical and cell biological experiments were carried out and data were analyzed by M.D.Z., C.M., S.A., S.E.R., A.J., P-A.L., D.F. and M.S. D.D. supervised S.A. and S.E.R. The paper was written by M.S. with contributions from other authors.

