## Supplemental Figures 1-4 for "The CIP2A-TOPBP1 complex safeguards chromosomal stability during mitosis"

**a**

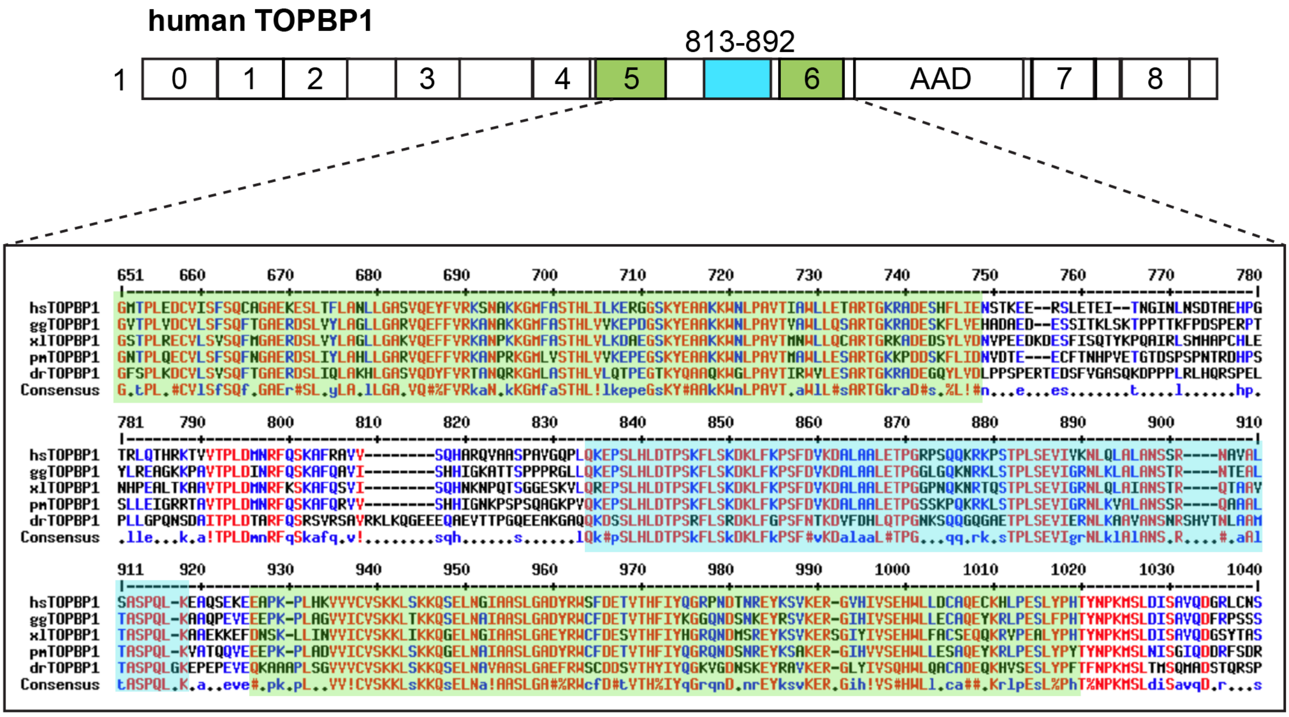


**b**

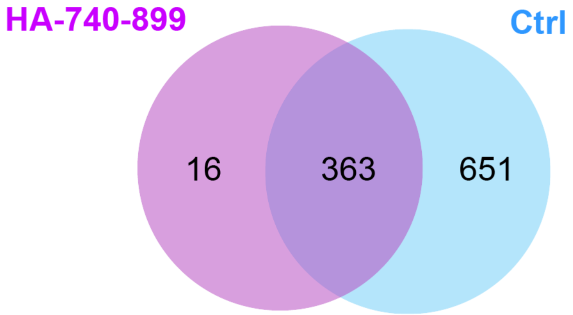


**c**

| Accession | Protein Name |
|---|---|
| Q92547 | DNA topoisomerase 2-binding protein |
| Q92598 | Heat shock protein 105 kDa |
| O95757 | Heat shock 70 kDa protein 4L |
| Q9NZL4 | Hsp70-binding protein 1 |
| O95816 | BAG family molecular chaperone regulator 2 |
| P12956 | X-ray repair cross-complementing protein 6 |
| Q96CS3 | FAS-associated factor 2 |
| Q9BSJ8 | Extended synaptotagmin-1 |
| Q15363 | Transmembrane emp24 domain-containing protein 2 |
| P28288 | ATP-binding cassette sub-family D member 3 |
| P13010 | X-ray repair cross-complementing protein 5 |
| Q8TCG1 | Protein CIP2A |
| A0A075B6Q5 | Immunoglobulin heavy variable 3 |
| O75600 | 2-amino-3-ketobutyrate coenzyme A ligase, mitochondrial |
| P54725 | UV excision repair protein RAD23 homolog A |
| Q86YZ3 | Hornerin |

**d**

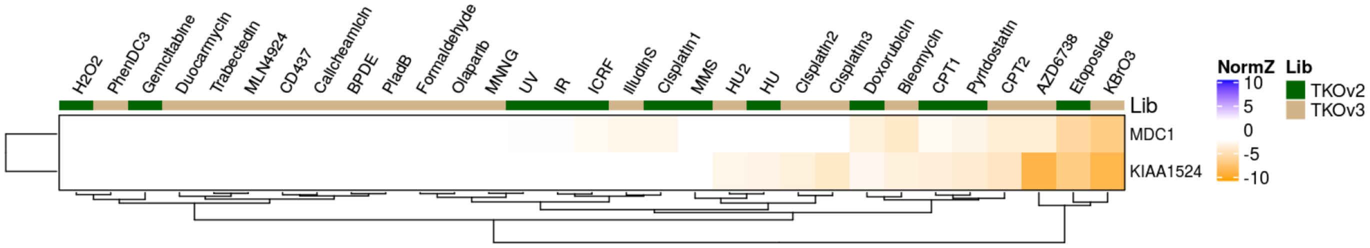


**Supplementary Fig. 1** Supporting data on TOPBP1 binds to CIP2A via a conserved interaction surface between BRCT5 and 6. **a** Sequence alignment of the region between BRCT5 and BRCT6 of TOPBP1. Examples from the five vertebrate families are included (hs: *homo sapiens*, mammalia; gg: *gallus gallus*, bird; xl: *xenopus laevis*, amphibia; pm: *podarcis muralis*, reptilia; dr: *danio rerio*, fish). Conserved amino acids are in red. **b** Venn diagram of the LC-MS/MS results. **c** List of proteins identified by LC-MS/MS specifically co-immunoprecipitated by HA-TOPBP1 amino acid 740-899, but not present in the control (HA-beads alone) Bait and protein of interest are highlighted in green and yellow, respectively. **d** Drug sensitivity correlation matrix between MDC1 and CIP2A (KIAA1524). Derived from https://durocher.shinyapps.io/GenotoxicScreens/

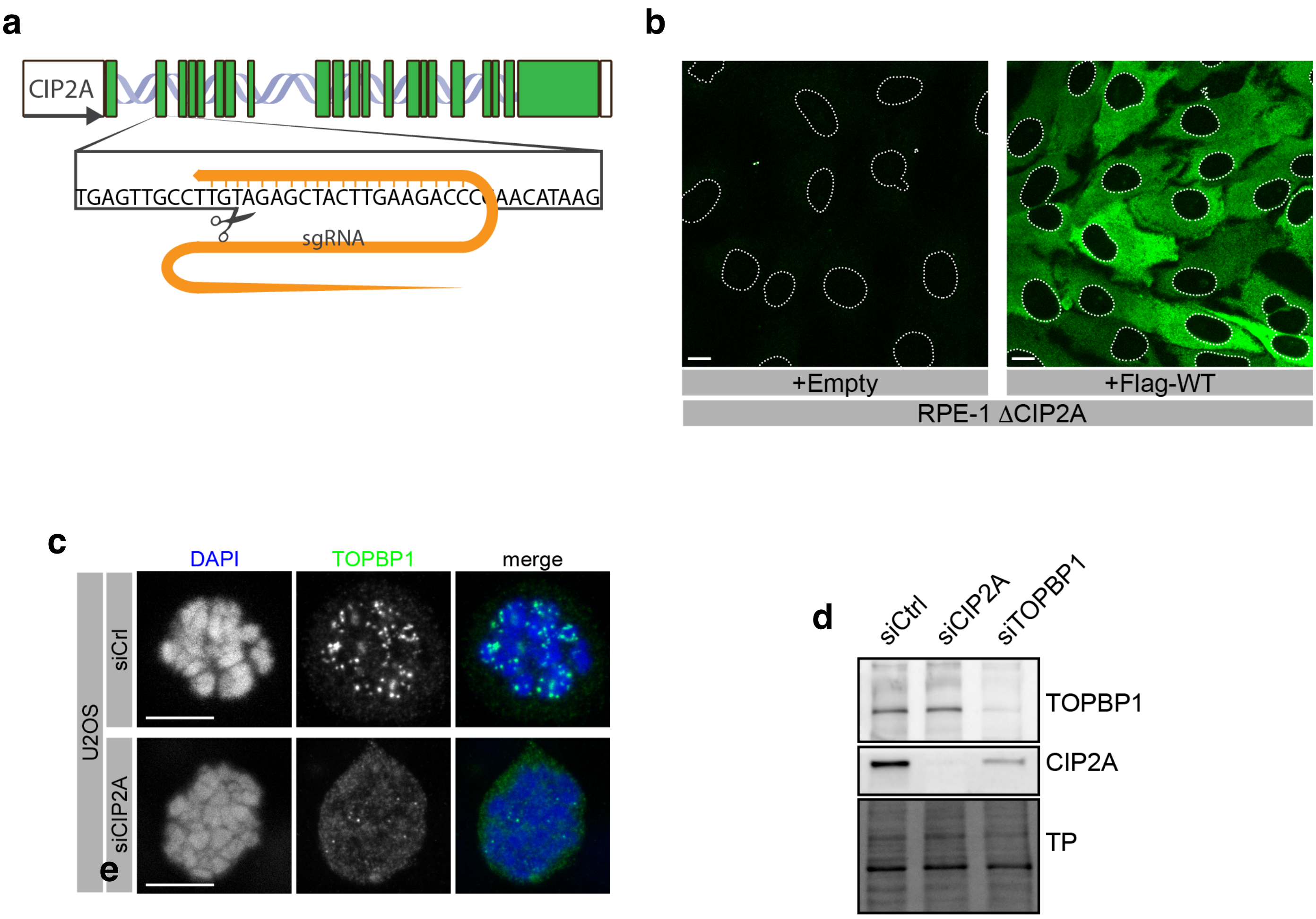


**Supplementary Fig. 2** Supporting data on CIP2A mediates recruitment of TOPBP1 to sites of DSBs in mitosis. **a** Schematic of the CIP2A knock-out strategy in RPE1 cells. **b** Confocal micrographs of untreated RPE1 ΔCIP2A cells, stably transduced with empty vector (+Empty) and with a lentiviral vector containing Flag-tagged CIP2A wild type cDNA (+Flag-WT) and stained for CIP2A. **c** Confocal micrographs (maximum intensity projection) of control siRNA (siCtrl) and CIP2A siRNA (siCIP2A) transfected, Nocodazole-arrested U2OS cells, 1h after irradiation with 1 Gy. **d** Western blots of total cell extract of U2OS cells, transfected with control siRNA (siCtrl), CIP2A siRNA (siCIP2A) and TOPBP1 siRNA (siTOPBP1).
All scalebars = 10 µm.

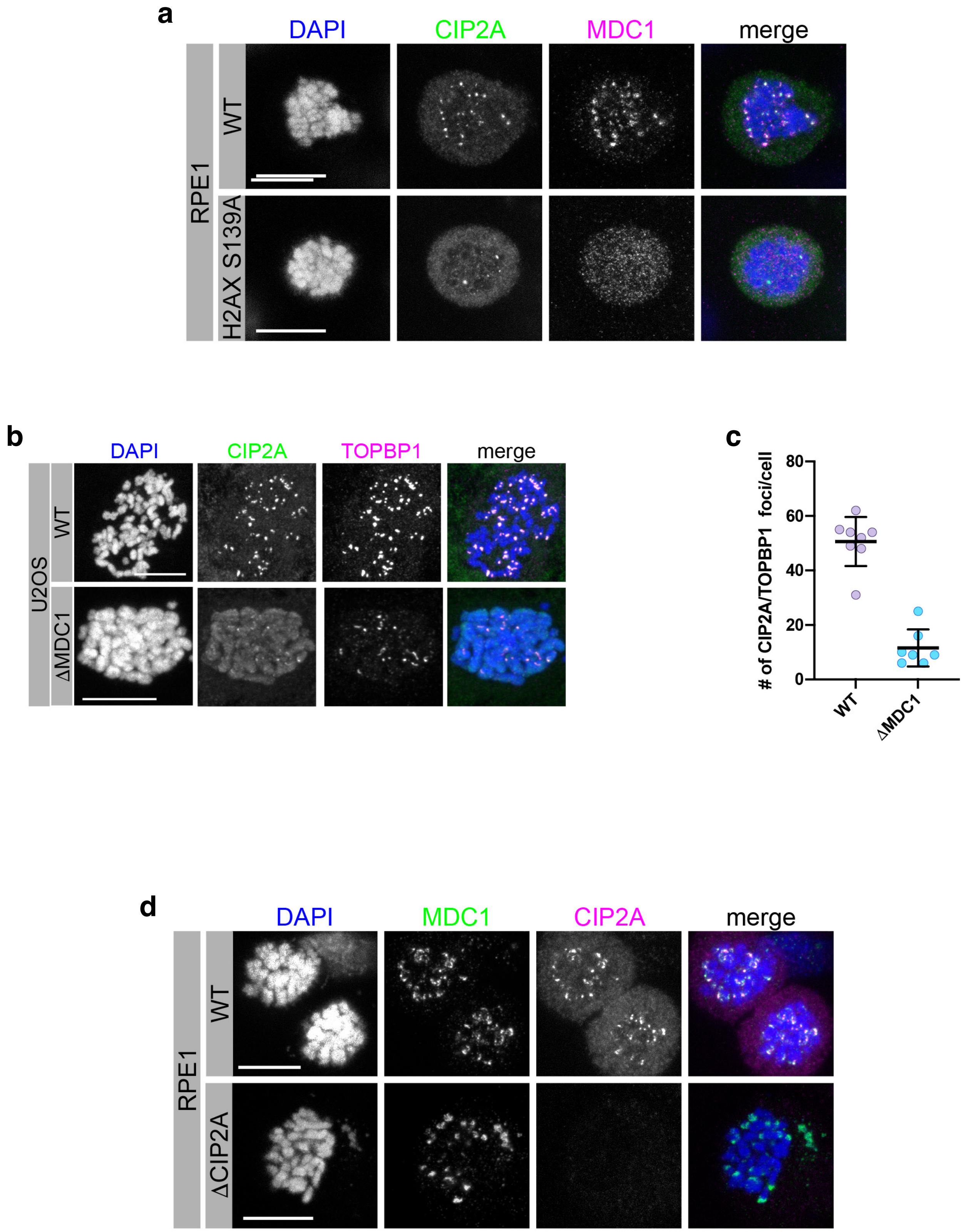


**Supplementary Fig. 3** Supporting data on MDC1-dependent recruitment of CIP2A-TOPBP1 to sites of mitotic DSBs. **a** Confocal micrographs (maximum intensity projections) of Nocodazole-arrested RPE-1 wild type and H2AX$^{S139A/S139A}$ knock-in cells, irradiated with 1 Gy and stained for TOPBP1 and CIP2A. **b** Confocal micrographs (maximum intensity projections) of Nocodazole-arrested U2OS wild type and U2OS ΔMDC1 cells, irradiated with 1 Gy and stained for TOPBP1 and CIP2A. **c** Quantification of the experiment in **b**; number of foci were manually counted; data points represent individual mitotic cells (n=7). **d** Confocal micrographs (maximum intensity projections) of Nocodazole-arrested RPE-1 wild type and ΔCIP2A cells, treated with 1 Gy and stained for MDC1 and CIP2A. All scalebars = 10 µm.

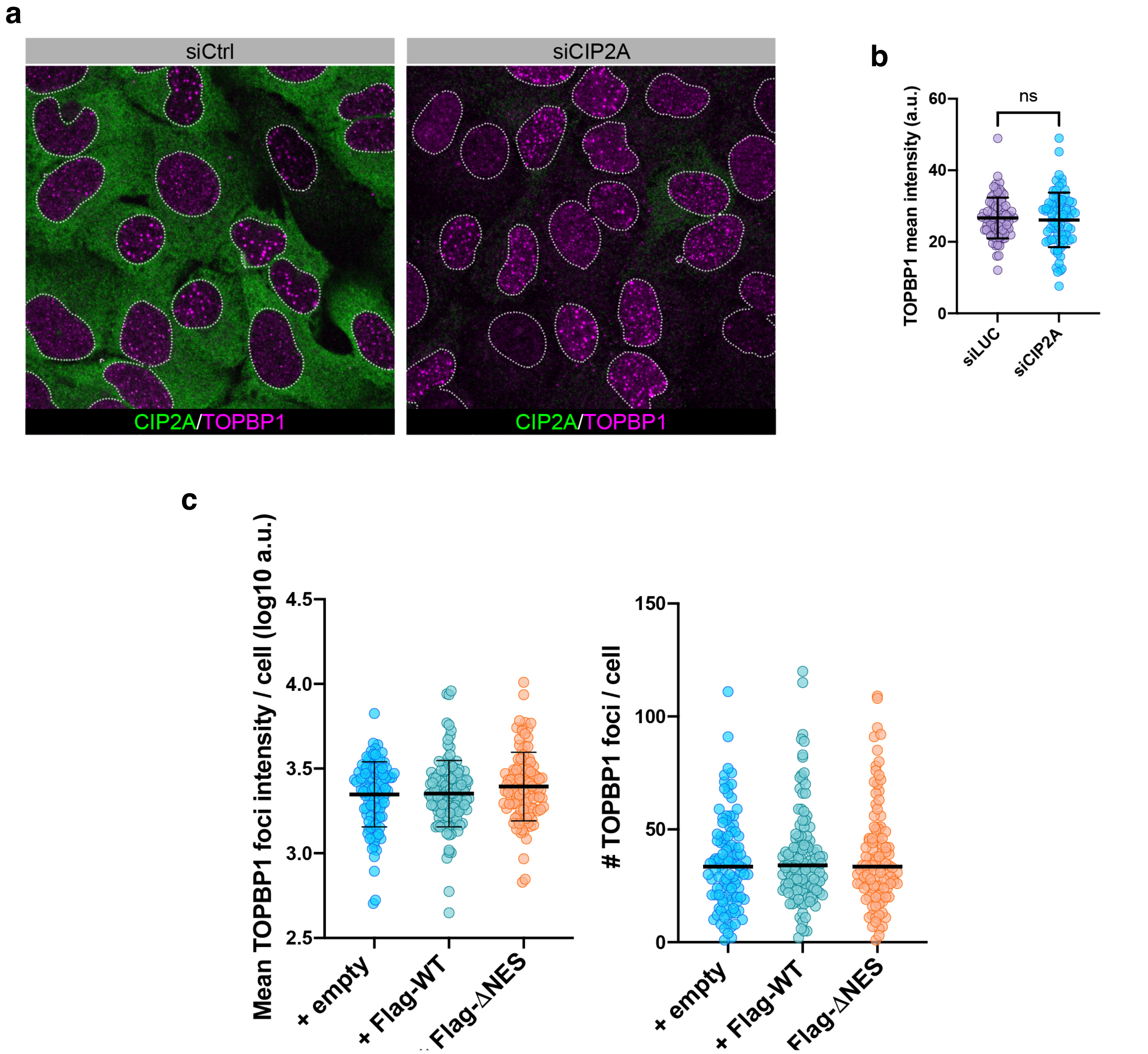


**Supplementary Fig. 4** Supporting data on CIP2A-independent TOPBP1 foci formation in interphase cells. **a** Confocal micrograph of unsynchronized U2OS cells, transfected with control siRNA (siCtrl) and CIP2A siRNA (siCIP2A), irradiated with 3 Gy and stained for CIP2A and TOPBP1. **b** Quantification of TOPBP1 foci mean intensity of the experiment in **a**. Statistical significance was calculated using unpaired t-test. Bars and error bars represent mean and SD; n=82 (siLUC), 74 (siCIP2A). **c** Quantification of TOPBP1 foci mean intensity and number of TOPBP1 foci per cell in interphase RPE1 ΔCIP2A cells stably transduced with empty vector (+empty) or Flag-tagged full-length CIP2A (+Flag-WT) and Flag-tagged CIP2A ΔNES (+Flag-ΔNES). Bars and error bars represent mean and SD.
